# Survival of Transplanted Retinal Ganglion Cell in Human Donor Eyes under Elevated Pressure

**DOI:** 10.1101/2025.04.22.650064

**Authors:** Shahna S. Hameed, Salil J. Gupta, Kathleen K. Ho, Tasneem P. Sharma

**Affiliations:** Eugene and Marilyn Glick Eye Institute, Department of Ophthalmology, Indiana University School of Medicine, Indianapolis, IN, 46202; Pharmacology and Toxicology, Indiana University School of Medicine, Indianapolis, IN, 46202; Stark Neurosciences Research Institute, Indianapolis, IN, 46202

**Keywords:** Glaucoma, cell therapy, transplantation, intraocular pressure, retinal ganglion cells

## Abstract

Glaucoma is a group of optic neuropathies characterized by visual field loss, classically due to increased intraocular pressure (IOP) and retinal ganglion cell (RGC) degeneration. Current treatment options reduce IOP, but progressive RGC degeneration persists. The ability to reprogram *de novo* RGCs from human corneal keratocytes provides a valuable tool to potentially restore vision in patients with late-stage disease when most RGCs are irreversibly damaged. We investigate the survival of these human induced pluripotent stem cell (hiPSC) derived RGCs after culturing them in human donor eyes under conditions of elevated and normal IOP using the pressurized ocular translaminar autonomous system (TAS) chamber. The hiPSCs were generated by reprogramming human donor keratocytes using Sendai viral vectors with Yamanaka factors. The hiPSCs were then differentiated into retinal organoids (ROs) from which RGCs were obtained. The RGCs were transduced with AAV2-CBA-EGFP (Adeno-Associated Virus serotype 2- Chicken Beta Actin- Enhanced Green Fluorescent Protein) and successfully transplanted into donor human eyes obtained from individuals having non-ocular history. They were pressurized for 5-7 days, with the left eye maintained at normal IOP and right eye at high IOP. Viability was measured by expression levels of pro-survival pathways via qRT-PCR, immunohistochemistry staining, and electroretinography (ERG) for retinal function. After RGC transplantation, increased expression of pro-survival and decreased inflammatory and apoptotic markers were identified in normal IOP conditions compared to high IOP. In conclusion, survival of RGCs was more conducive under normal IOP conditions with significantly increased degeneration observed at high IOP. Thus, suggesting that elevated IOP could potentially create a microenvironment that would significantly inhibit successful transplantation of *de novo* RGCs.

## 1. INTRODUCTION

Glaucoma is a group of optic nerve diseases caused by increased IOP which leads to optic nerve degeneration and RGC death, ultimately causing permanent irreversible vision loss (Kwon *et al*., 2009). The RGCs are the neurons that are specifically affected in glaucoma and transmit information to visual centers of the brain. ^12^ Since RGCs are terminally differentiated central nervous system neurons, they cannot heal once damaged (Bagnis *et al*., 2011).

Elevated IOP, evident in most glaucoma patients, is the only modifiable major risk factor for glaucoma. The current treatment options for the disease aim to delay disease progression by lowering IOP (Kass *et al*., 2002). However, this strategy does not control progression in all glaucoma patients and there is no current treatment that can regenerate the damaged RGCs at late-stage glaucoma (Aubert *et al*., 1995; Nicholls and Saunders, 1996; Di Giovanni *et al*., 2005). Transplantation of RGCs presents a viable therapeutic strategy, however, transplantation efficiency remains at 1% as observed by previous studies (Venugopalan *et al*., 2016; Wu *et al*., 2018). Effective transplantation could be hindered by several variables, including cell density, cell aggregation and shearing during transplantation, implications of the microenvironment, integration efficiency, or local and systemic immune responses (Abud *et al*., 2022; Zhang *et al*., 2021)

Cellular replacement strategies include primary RGCs, embryonic stem cell derived neural progenitors, Mueller glia derived precursors, and pluripotent stem cells (Soucy *et al*., 2023). Most transplantation strategies have been tested in animal model systems and while rodent models do not precisely mimic the human retina, there have been studies of transplantation of mouse and human RGCs into a mouse model (Croteau *et al*., 2022; Oswald *et al*., 2021; Vrathasha *et al*., 2022; Wu *et al*., 2021). Previously, it has been shown that mouse induced pluripotent stem cell (iPSC) derived RGCs had a 65% success rate with axon growth extending into the optic nerve head within rodent host retinas (Oswald *et al*., 2021). Further, it has been shown that human RGC growth is supported by neurotrophic factors in mouse retina explants (Croteau *et al*., 2022). Another study identified that mouse embryonic stem cells integrate and form synapses in an RGC depleted mouse model (Wu *et al*., 2021). There is also evidence that transplanted embryonic stem cell-derived neural progenitors improve visual function in mice with depleted RGCs (Divya *et al*., 2017). Furthermore, it has been shown that transplanted human iPSC derived RGCs were electro-physiologically functional in living mouse retinas (Vrathasha *et al*., 2022). Still, survival of these cells is highly limited, with only a few percent surviving after 5 months which could potentially be explained by lack of immune suppression in the mice and the use of human xenograft transplantation (Oswald *et al*., 2021). Cell survival is crucial for advancement of these cutting-edge techniques and understanding methodologies to increase survivability of transplanted cells would allow us to improve future cell therapeutic strategies.

To optimize transplantation methodology, we aimed to study RGC transplantation using *de novo* RGCs derived from reprogramming corneal fibroblasts of human donor eyes. Using human tissues and cells also allows us to consider demographics, so our human eye model system allows us to increase applicability to human eyes, and the chance of bringing our discoveries from bench to bedside. Our approach of a pre-clinical human model system to test transplantation efficiency reduces the use of animal models in research, consistent with the FDA’s Alternative Methods Working Group recommendations to improve clinical translatability (Avila *et al*., 2020). While rodent models provide valuable insights into physiological responses, safety, efficacy, and other clinical evaluations, certain morphological differences between their ocular structures and human architecture may impact their translational efficacy (Nucci et al., 2018; Avila et al., 2020). Notably, mice, rats, and rabbits either lack a well-defined lamina cribrosa or possess an underdeveloped one (Burgoyne, 2015; Raghunathan *et al*., 2017). Furthermore, differences in cellular composition, particularly in collagen types, exist among these species (Albrecht May, 2008). Therefore, modeling glaucoma using human eyes will further add to our knowledge of the disease.

Human iPSCs offer the potential of generating de novo RGCs and others are studying their use in autologous transplantation. Specifically, autologous iPSC derived retinal pigmented epithelial cells have been studied in numerous human clinical trials with limited success (Raimondi *et al*., 2022). A clinical trial performed for dry age related macular degeneration (Mandai *et al*., 2017) depicted discrepant results between patients. There is an in-progress clinical trial for wet age-related macular degeneration using retinal pigment epithelium transplantation (Clinicaltrials.gov ID: NCT04339764) (2020) and several other trials at various stages of completion using different stem cell origins and cell types (Feldman, 2023), however, no transplantation trials have been conducted in humans using RGCs. The goal of our study was to utilize RGCs for transplantation in human donor eyes in an *ex vivo* model.

To effectively understand downstream transplantation effects, there is a need to study RGC survival in the glaucomatous retinal microenvironment (Liu and Lee, 2021). Even though glaucoma pathogenesis is being controlled with IOP-lowering therapeutics and surgeries, other options are currently being explored that target the RGCs directly and include neuroprotection, gene therapy, and cell replacement. A critical question is the survival of the RGCs within the human retina for therapeutic success. For a beneficial impact, RGCs must integrate within the retina through their interneurons and their axons need to make the correct connections to the visual centers of the brain. But, before this can be explored, we must investigate the survival of these cells in the human retina, which is the primary goal of this work. From previous literature, we know that the survival rate of RGCs in animal models after transplantation is typically less than 1% (Chao *et al*., 2017; Venugopalan *et al*., 2016). Factors mediating survival include trophic support, graft rejection, reactive gliosis, optic neuritis, and oxygen and nutrient supply (Zhang *et al*., 2021). Nevertheless, no one has studied the intricacies of transplantation within human eyes under conditions of pressure or axotomy. It is crucial to study the survival of RGCs in human donor eyes to understand how the microenvironment can impact effective transplantation. To this end, we studied survival of RGCs after transplantation in axotomized human eyes under elevated versus normal pressure conditions.

## 2. MATERIALS AND METHODS

### 2.1. Human donor tissue collection and demographics

The human ocular tissues were obtained from Lions Gift of Sight Minnesota eye bank and Lions World Vision Institute in Tampa, Florida. They were then processed according to the Declaration of Helsinki. The methods were performed in accordance with the relevant guidelines and regulations, approved by the Indiana University School of Medicine on Non-Human Subject Research and in accordance with the guidelines of Association of Research in Vision and Ophthalmology (ARVO) for handling de-identified specimens. Experiments for human posterior segment culture experiments were compliant with the approved Indiana University - Institutional Biosafety Committee protocol (Proposal # IBC 1318). All the tissues perfused were from subjects having no ocular history or any other central nervous system pathology. All experiments were performed in six independent repeats.

### 2.2. Generation of *de novo* RGCs

The iPSCs derived from human keratocytes were differentiated into RGCs (Hameed and Sharma, 2025). The iPSCs were generated using CytoTune^TM^ iPSC 2.0 Sendai reprogramming kit (Thermo Fisher Scientific, Cat. No. A16517, Lot no. L2190071), which is based on three non-integrative Sendai viral vectors containing hKOS, hc-Myc and hKlf4 as per manufacturers’ instruction. Karyotyping was performed to assess the chromosomal loss during reprogramming (Hameed and Sharma, 2025). The generated iPSCs were differentiated into ROs as per previously published protocol with a slight modification (Sharma *et al*., 2017) (Hameed and Sharma, 2025). In brief, iPSCs were dislodged from the plate and dissociated into single cell suspension using TryPLE Express enzyme (Thermo Fisher scientific, Cat. No. 12604-013). The iPSCs were seeded at a density of 1.0 × 10^4^ cells per will into an ultra-low adhesion 96-well plate (Corning Costar, Cat.no. 7007) with 3D differentiating media with 20mM ROCK inhibitor (Millipore, Sigma, Cat. No. 6880001-1MG) and 3nM IWRe (Cayman Chemicals, Cat. No. 13659). After 24 hours, 1% extracellular matrix (ECM) (Matrigel) (Corning, Cat. No. 354230) was added to the media. Every 48 hours 50% of the media was replaced with fresh media and on day 12, ROs from all 96wells were carefully transferred to a 10cm^2^ dish (Greiner Bio-one, Cat. No. 664970). The organoids were fed with new 3D differentiating media containing 1% ECM. From days 15-18, the ROs were fed with 3D differentiating media with1% ECM, 3 mM CHIR (Millipore Sigma, Cat. No. SML1094-5MG) and 100nM SAG (Millipore Sigma, SML1314-1MG). On day 18, 3D differentiating media was switched to neural retinal culture medium (DMEM F12 – Thermo Fisher Scientific, Cat. No.10565018; 2mM Glutamax-Thermo Fisher Scientific, Cat. No. 35050061; N2 supplements – Thermo Fisher Scientific, Cat. No. 17502-048; Primocin - Invivogen, Cat. No. ant-pm-1) and continued until day 30. The day 30 organoids were used for further downstream generation of RGCs. In brief, 10 ROs were used for enzymatic dissociation with papain for 5 minutes at 37°C. The dissociated cells were collected by brief centrifugation at 800 rpm for 3 minutes. The cells were dissociated in complete neurobasal media prepared as previously published (Hameed *et al*., 2024b). The cells were seeded into an 8 well chamber slide coated with Poly-D-Lysine and laminin at a density of 2.0 × 10^4^ cells/ well.

### 2.3 Characterization of keratocytes, their derived iPSCs and RGCs

Keratocytes, ROs containing RGCs, and dissociated RGCs were characterized using immunofluorescence. For iPSCs, pluripotency markers were assessed using both immunofluorescence and PCR (*SOX2, KLF4, NANOG, C-MYC*) (Hameed and Sharma, 2025). For immunostaining, the cells or ROs after the sectioning were processed as follows. In brief, after washing with 1X PBS, the cells and RO sections were fixed in 4% formaldehyde fixation for 10 minutes at room temperature. After washing thrice with 1X PBS, the cells or sections were permeabilized in 0.5% Triton X-100 for 20 minutes followed by blocking using SuperBlock^TM^ T20 (Thermo Fisher Scientific) for 1 hour. The respective primary antibodies (Keratocytes: Keratocan, Thermo Fisher Scientific, Cat. No. BS-11054R; iPSCs: TRA-1-60, Abcam, Cat. No. AB16288 and SOX2 (Abcam, Cat. no. ab97959, RGCs: ISLET 1, Abcam, Cat. No. AB109517, THY1: Abcam, Cat. No. AB181469, RBPMS, Abcam, Cat. no. ab152101 and NOVUS, Cat. no. NBP2-03905, BRN3A, EMD Millipore, Cat. No. MAB1585) were diluted in blocking buffer and incubated overnight at 4^°^C. After washing primary antibodies thrice with 1X PBS, the cells or RO sections were incubated with their respective secondary antibodies (Goat anti-Rabbit IgG, Alexa Flour 568, Invitrogen, Cat. No. A11036, Goat anti-Mouse IgG, Alexa Flour 488, Invitrogen, Cat. No. A32723, Goat anti-Mouse IgG, Alexa Flour 568, Cat. No. A10037) for 2 hours at room temperature. After washing thrice with 1X PBS, cells or RO sections were mounted using ProLong^TM^ Gold antifade reagent with DAPI (Thermo Fisher Scientific) and visualized under a fluorescent microscope at 20X magnification and images captured.

### 2.3. Transduction of hiPSC derived RGCs with AAV2-CBA-EGFP

The RGCs dissociated from day 30 ROs at a density of 2.0X10^4^ were transduced with AAV2-CBA-EGFP (Vector Biolabs, Cat. No. 7072, Lot No. A11M1031, Titer: 1X10^10^ viral genomes/mL). The viral media was removed 16-18 hours of post-transduction, and the cells were maintained in RGC culture media for another 72 hours. After confirmation of successful transduction by assessing the GFP expression via fluorescent microscopy, and viability of the transduced cells being transplanted through trypan blue cell viability assay, ∼1.0 × 10^4^ of transduced cells, were isolated and seeded into the donor posterior segments.

### 2.4. Seeding of hiPSC-AAV2-CBA-EGFP tagged RGCs into donor posterior segments with pressurized perfusion

Human donor eyes harvested 6–12 h post-mortem were received within 24 hours within a moist chamber and shipped at 4 °C in a sealed container to be utilized for experimental analyses. The donor eyes were dissected as previously described (Peng *et al*., 2022; Sharma *et al*., 2020). In brief, the extra-ocular donor eye surface was washed thrice with phosphate buffered saline (PBS) and then dissected cross-sectionally at the equator into anterior and posterior segments. The lens and the vitreous humor were removed from the posterior eyecup without disturbing the retina. For transplantation, the transduced RGCs after 72 hours of AAV2-CBA-EGFP viral transduction were collected for seeding. For harvest from tissue culture plate, the RGCs were trypsin digested for 3 minutes. The trypsin activity was then arrested with complete neurobasal media prepared as previously published (Sharma *et al*., 2015; Pang *et al*., 2007; Peng *et al*., 2022; Hameed *et al*., 2024a). Briefly, the complete neurobasal medium was prepared utilizing penicillin (100 U/mL, Cytiva), GlutaMAX (2 mM, Thermo Fisher Scientific), streptomycin (100 µg/mL, Cytiva), pyruvate (1 mM, Thermo Fisher Scientific), insulin (5 µg/mL, Gibco), transferrin (100 µg/mL, Gibco), sodium selenite (40 ng/mL, Gibco), thyroxine (100 ng/mL, MilliporeSigma), tri-iodothyronine (40 ng/mL, MilliporeSigma), progesterone (60 ng/mL, MilliporeSigma), putrescin (16 µg/mL, MilliporeSigma), forskolin (5 µM/mL, MilliporeSigma) and 1% human serum (Gibco) with basal neurobasal medium (Gibco) containing 10X B27 supplements (Gibco).

The RGCs were collected after centrifugation at 800 rpm for 5 minutes. The supernatant was removed, and the RGCs were gently mixed with 100μL of Matrigel (Corning, Cat. No. 354230, growth factor reduced) and this mixture was applied to the retina within the posterior segments. To enhance the stability of the mixture on the retinal surface, a soft contact lens (Acuvue) with 1 diopter power was gently placed over it. After 2 hours of seeding, 2 mL of complete neurobasal media was added on top of the contact lens. The eye cup including sclera and the optic nerves were also bathed in complete medium to avoid drying out of the tissue. The eyes were then incubated at 37 °C in a CO_₂_ incubator under standard cell culture conditions with maintained humidity for 24 hours. Following incubation, the dissected cups with seeded RGCs were then placed within the TAS model and perfused as described previously (Peng *et al*., 2022; Sharma *et al*., 2020). In brief, the posterior cup was carefully placed in the dome of the IOP chamber of the TAS model with the optic nerve facing upwards. The IOP chamber was tightly sealed with an epoxy resin O ring with four screws. The ICP chamber, with the optic nerve inserted, was precisely positioned over the IOP chamber and securely sealed. The in-flow and out-flow tubing were inserted into the respective portals of the IOP and ICP chamber. The inflow syringes were filled with complete neurobasal perfusion media and connected to an automated pump with an adjusted infusion flow rate of 0.3-0.5 µL/minute for attaining normal IOP and 1-2 µL/minute for attaining high IOP. The outflow syringes were connected to manually calibrated hydrostatic pressure transducers attached to a multichannel bridge amplifier. The OD eye (oculus dexter, right eye) was perfused at high IOP condition, and the paired OS eye (oculus sinister, left eye) was perfused at normal IOP condition. The pressure data were recorded every minute and the average IOP for every 24 hours was calculated with the LabChart software system. The perfused eyes were incubated at 37°C in a CO_₂_ incubator for 6 days under standard cell culture conditions with maintained humidity for 24 hours, after which the tissues were harvested for downstream experiments.

### 2.5. Quantitative gene expression analysis

RNA extraction of pre and post retinal tissues were performed using MACHEREY-NAGEL kit and was conducted as per the manufacturer’s protocol. For pre-perfusion, 2 mm size peripheral retinas harvested from the OD (N=6) and OS (N=6) eyes prior RGC seeding and perfusion (N=6). A total of 100 ng of RNA was reverse transcribed into cDNA using iScript^TM^ kit (Bio-Rad Laboratories) as per manufacturer’s protocol. To assess the suitability and viability of the donor retinas for use in perfusion culture, the cDNA obtained from pre-perfusion eyes were subjected for quantifications of inflammatory (*TLR4, GFAP, GSS, AIF1*), apoptosis (*BAX, CASP3, CASP7*) and retinal (*RBPMS, ISL-1, THY1, NEFH, RHO, PRKCA*) markers. To evaluate the RGC survival and apoptosis, cDNA obtained from the post-perfusion retinas were assessed for the genes of apoptosis (*BAX, CASP3)* and RGC marker (*THY1*). Further, to evaluate whether the apoptotic and inflammatory genes is derived from donor cells or host cells, we compared the expression of apoptotic (*BAX, CASP3, CASP7*), and inflammatory genes (*TLR4*) between eyes perfused at normal IOP without transplantation (N=3) and those perfused at normal IOP with transplanted RGCs (N=6). Likewise, we also compared the gene expression in eyes perfused at high IOP without transplantation (N=3) with those with transplanted RGCs (N=6). TaqMan array qRT PCR was used for assessing the pre and post perfusion retinal gene expressions. The reaction mix was composed of TaqMan fast advanced master mix (Thermo Fisher Scientific, Cat. No. 4444558), cDNA and primers for genes of interest coated in a pre-designed TaqMan array plate. The qRT PCR was performed on an Applied biosystems QuantStudio3 system for a total reaction volume of 10 µL/sample. For all the gene expression analysis, the fold change (FC) was calculated using the 2^-ΔΔct^ method by keeping GAPDH as a housekeeping gene.

### 2.6. Immunostaining of the retinal flat mounts

Immunostaining for the RGC marker RBPMs was performed on RGC-transplanted retinas that were perfused under both normal and high IOP conditions. In brief, post perfusion, the retinas were collected from the RGC transplanted eyes and fixed in 4% formaldehyde (Spectrum Chemical MFG Corp, Cat no. F-262) for 15 minutes. After washing with PBS, the sections were permeabilized and blocked for 45 minutes in blocking buffer containing Tween 20 (SuperBlock^TM^ T20, Thermo Fisher Scientific, Cat. No. YK378727). The retinal sections were incubated overnight at 4°C with mouse RBPMS primary antibody (Novus, Cat. No. NBP2-03905, 1:300). Followed by washing with PBS, the retinal flat mounts were stained with its secondary antibody (Goat anti-Mouse IgG, Alexa Flour 568, Cat. No. A-11004, 1:300) for 1 hour at room temperature. After PBS wash, tissues were mounted using ProLongTM Gold antifade reagent with DAPI (Thermo Fisher Scientific, Cat. No. P36935) and visualized under a fluorescent microscopy at 20X magnification.

### 2.7. Sectioning and staining of the optic nerve head

Optic nerve heads were collected from RGC-transplanted eyes after perfusion at both normal and high IOP. The optic nerves were embedded in Optimal Cutting Temperature (O.C.T) compound (Fisher healthcare, Cat. No. 4585) and stored in -80°C overnight. Using a cryostat (Leica), 30 microns thick sections of tissues were taken and used for assessing the markers ECM deposition. In brief, the sections on the slides were heated for 45 minutes at 60°C to remove the O.C.T compound. The adhered sections were then washed with PBS and fixed in 4% formaldehyde for 15 minutes. Following PBS washing, the sections were permeabilized and blocked in blocking buffer containing tween 20 (SuperBlock™ T20, Thermo Fisher Scientific, Cat. No. YK378727). The sections incubated at 4°C with antibodies such as collagen IV (Novus, Cat. No. NB120-6586, 1:1000), fibronectin (Millipore, Cat. No. AB 1945, 1:1000) and laminin (Novus, Cat. No. NB300-144, 1:1000). The sections were washed with PBS and stained with the secondary antibody (Goat anti-Rabbit IgG, Alexa Flour 568, Cat. No. A-11011, 1:300). After three PBS washes, the tissues were mounted using ProLong™ Gold Antifade Reagent with DAPI (Thermo Fisher Scientific, Cat. No. P36935). Images were captured under a fluorescent microscope at 4X magnification. Mean fluorescent intensity was calculated using ImageJ, and background signals were subtracted with a secondary antibody control.

### 2.8. *Ex vivo* electroretinography (ERG) for measuring retinal function

The *ex vivo* ERG was performed as previously published (Hameed *et al*., 2024b). In brief, 2mm diameter central retinal sections were collected from perfused eyes in neurobasal medium. Prior to ERG evaluation the sections were dark adapted for 15 minutes. The tissues were carefully divided into two equal halves and positioned onto the OcuScience *ex vivo* sample holder with the photoreceptor layer upwards. The OcuScience ERG system parameters such as Impedance (≤ 20KOhms), temperature (37°C) and offset voltage (≤ 10mV) were adjusted. The Ames’ medium of pH 7.4 was prepared and bubbled to control O_2_ and CO_2._ Tissue samples were perfused with Ames′ Medium buffer at 50 mL/hour, and full field flash ERG recording was performed using handheld multi-species ERG unit (HMsERG by OcuScience). The ERG responses were recorded by stimulating the retina with sequential flashlights with varying intensities from 1mcd flash intensity to 1000 mcd. The amplitudes were measured from baseline to the positive peak of each wave form. Implicit time was measured by time to peak of the corresponding a-wave and b-wave component. The b/a ratio, a and b-wave implicit time at a low flash intensity of 30 mcd were calculated. Data were analyzed between the RGC transplanted eyes at high IOP versus transplanted eyes at normal IOP.

### 2.9. Statistical Analysis

GraphPad Prism software was used to analyze all the results. We quantitatively assessed gene expression of donor retinas at pre-perfusion stage via two-way ANOVA. To assess the significance of differences in pressure perfusion between transplanted eyes at high and normal IOP, we performed two-way ANOVA and student’s t-test. Two-way ANOVA was performed to calculate the significance of genes of apoptosis and inflammation and also for assessing the significance of ECM deposition. Student’s t-test was done for assessing *THY1* expression. All the ERG data was analyzed using Student’s t-test. In all experiments *p*<0.05 was considered statistically significant. The data has been presented as Mean ± SEM, the out layers removed by prism. All the experiments were done in six independent repeats.

## 3. RESULTS

### 3.1. Demographics of study subjects

The detailed demographics of the donor eyes used for the study have been provided in **Table 1**. All the donors were Caucasians. For the transplantation and pressure-perfusion studies, we used four males and two female donors. The medical history of all donor eyes was evaluated, and no ocular history was found to be associated with any of the donors. The average age of the donor used for the transplantation and perfusion experiment was 75.8 ± 5.4 years.

**Table 1.**
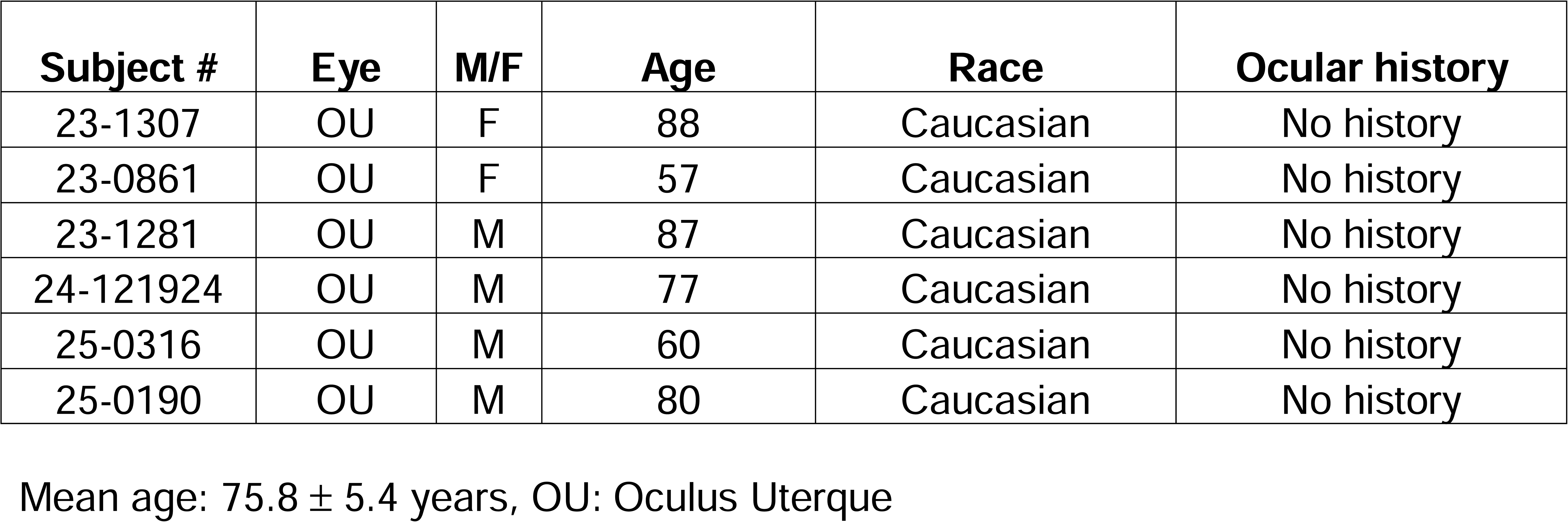
Detailed demographics of donor eyes used for RGC transplantation perfusion studies.

| **Subject #** | **Eye** | **M/F** | **Age** | **Race** | **Ocular history** |
| --- | --- | --- | --- | --- | --- |
| 23-1307 | OU | F | 88 | Caucasian | No history |
| 23-0861 | OU | F | 57 | Caucasian | No history |
| 23-1281 | OU | M | 87 | Caucasian | No history |
| 24-121924 | OU | M | 77 | Caucasian | No history |
| 25-0316 | OU | M | 60 | Caucasian | No history |
| 25-0190 | OU | M | 80 | Caucasian | No history |

Mean age: $75.8 \pm 5.4$ years, OU: Oculus Uterque

### 3.2. Successful generation of RGCs from human donor keratocytes

To evaluate the feasibility of RGC transplantation in the human eye, we utilized RGCs derived from donor keratocytes through iPSC technology. The schematic of the experimental paradigm has been provided in Fig. 1A. Control donor keratocytes were successfully cultured in the lab from a 77-year-old Caucasian male donor with no known ocular history. The cells were confirmed as keratocytes by immunofluorescence staining with the marker keratocan (Fig. 1B). Following this, reprogramming was performed on keratocytes for conversion to iPSCs using the Sendai virus-based reprogramming technology. The resulting iPSCs were characterized by immunofluorescent with expression of pluripotency marker TRA-1-60 (Fig.1C), SOX2 (Fig. 1D), and also by PCR with the expression of markers such as *C-MYC, SOX2, KLF4* and *NANOG* (Fig. 1E). These iPSCs were karyotyped which ensured no chromosomal loss during reprogramming (Fig.1F). The karyotyped iPSCs were further used to generate ROs (Fig.1G). The *de novo* generation of RGC in the ROs depicted positive staining for RGC markers such as ISLET 1, THY1 (Fig 1H) RBPMS and BRN3A (Fig. 1I). The nuclear specificity of BRN3A was confirmed using secondary antibody control (Supplementary Fig.1). To further isolate and transplant these RGCs, we performed papain based enzymatic digestion of ROs at day 30. The resulting RGCs were then cultured on poly-D-lysine and laminin dishes (Fig. 1J). Positive staining for RBPMS, BRN3A and THY1 was observed in the 2D cultured RGCs (Fig.1 K-M) validating the identity of RGCs.

**Fig. 1.**
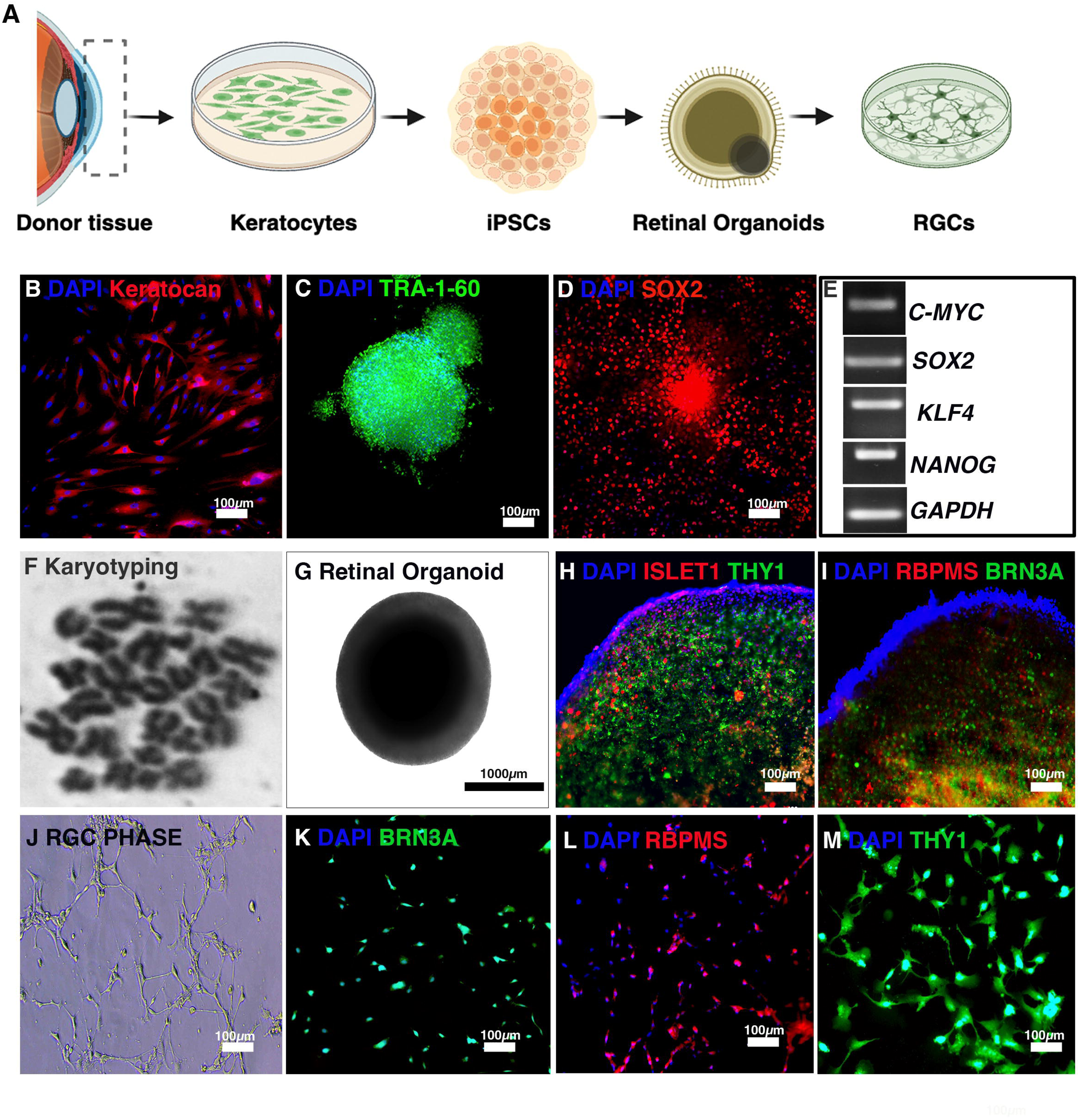
Generation of RGCs from human donor keratocytes. A). Illustrating the experimental paradigm used to generate human iPSC derived RGCs. Identification of positive marker expression of (B) keratocytes by keratocan and (C) iPSCs confirmed by TRA-1-60 expression, and (D) SOX2 expression. E) PCR analysis of pluripotency markers, including C-MYC, SOX2, KLF4, and NANOG, with GAPDH as the normalization control. F) Karyotyping of iPSCs depicting no chromosomal loss, (G) phase contrast image showing iPSCs were differentiated into ROs (retinal organoids) H) Immunostaining of ROs confirming RGC identity with positive expression of ISLET1, THY1, and (I) RBPMS, BRN3A. (J) Dissociation of ROs into single cells, followed by immunostaining confirming RGC identity through the expression of (K) BRN3A, (L) RBPMS, and (M) THY1. TRA-1-60: T cell receptor alpha locus associated 1-60, *SOX2*: SRY-box 2, *C-MYC*: cellular Myc, *KLF4*: Kruppel-like factor 4, *NANOG*: Nanog homeobox, *GAPDH*: glyceraldehyde-3-phosphate dehydrogenase, ISLET-1: Insulin gene enhancer protein 1, THY1: thymus cell antigen 1, iPSC: Induced pluripotent stem cells, ROs: Retinal organoids, RGC: Retinal ganglion cells, Magnification 20X

### 3.3. Successful transplantation of AAV2-CBA-EGFP tagged RGCs into human donor eyes

To minimize the influence of genetic and gender-related biases on RGC transplantation and to evaluate donor retinal viability, we assessed expression of retinal, apoptotic and inflammatory genes in these eyes via TaqMan arrays (Fig. 2) prior to RGC transplantation and pressure perfusion. All eyes used in the RGC transplantation underwent this initial evaluation phase. The gene expression data clearly showed measurable expression of retinal genes with non-significant difference in any of the genes between OD and OS eyes (*RBPMS, ISL-1, THY1, NEFH, RHO, PRKCA, STXA1)* prior to perfusion (Fig. 2A). Moreover, we did not find any significant changes in genes of apoptosis (*BAX, CASP3, CASP7*) (Fig. 2B) inflammation and gliosis (*TLR4, GFAP, GSS, AIF1*) (Fig. 2C), in the donor eyes prior to perfusion. After confirming viability through the expression of retinal markers and validating other gene markers in the donor eyes at the pre-perfusion stage, we proceeded with RGC transplantation in these eyes, followed by pressure perfusion within the TAS model as illustrated (Fig. 3A). The RGCs were seeded at a density of 1 × 10^4 cells per 2D culture dish for subsequent transplantation experiments. To verify the identity of the RGCs, they were stained with the specific RGC marker BRN3A. As depicted in Fig. 3B, RGCs derived from ROs in the 2D culture exhibited positive staining for BRN3A, confirming their RGC identity. To distinguish transplanted RGCs from native RGCs in the donor retinas, the *de novo* RGCs were transduced with AAV2-CBA-EGFP. Transduced RGCs showed GFP expression 48 hours post-transduction, with fluorescence observed after 72 hours post-transduction (Fig. 3C). Prior to seeding of the transduced RGCs, the viability of the cells was estimated using trypan blue cell viability assay and it showed only a small percentage of cell death in the transduced RGCs (Live cells: 77.64 ±5.6% Dead Cells: 22.3 ± 5.6%, N=4, Fig. 3D). Isolated GFP positive RGCs were collected and layered with 100 µL of Matrigel to be subsequently seeded onto the retina within the dissected posterior segment.

**Fig. 2.**
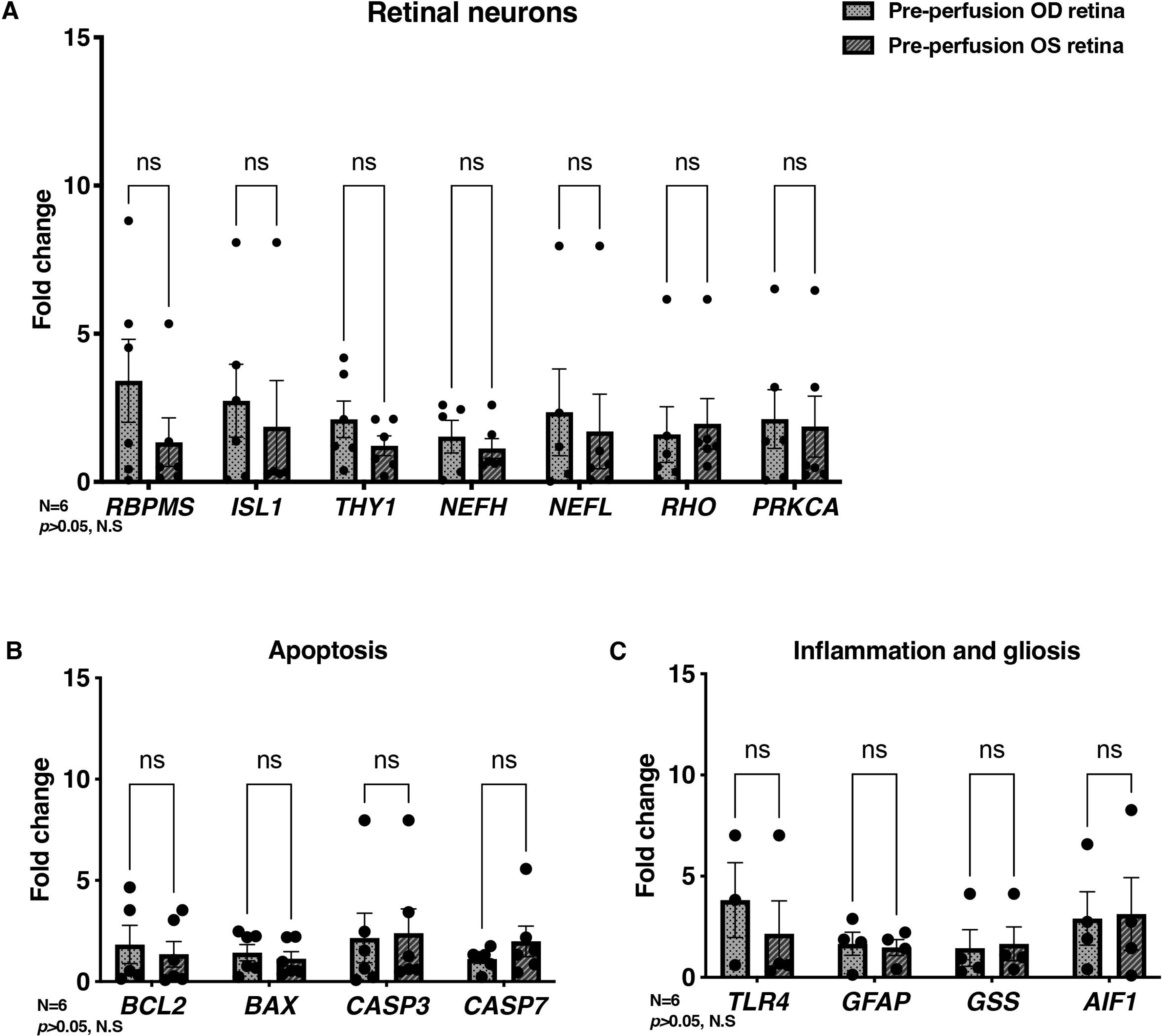
Confirmation of gene expression within the pre-perfused human donor eyes. A) The markers of retinal neurons (*RBPMS, ISL1, THY1, NEFH, NEFL, RHO, PRKCA*), (B) apoptosis (*BCL2, CASP3, CASP7*) and (C) inflammation and gliosis (*GFAP, GSS, AIF1, TLR4*) in the donor eyes assessed via quantitative real time PCR. *RBPMS*: RNA binding protein with multiple splicing*, THY1*: Thymus cell antigen 1*, ISL1*: Insulin gene enhancer protein 1*, NEFH*: Neurofilament heavy chain*, NEFL*: Neurofilament light chain, *RHO*: Rhodopsin*, PRKCA:* protein kinase C alpha, OD: Oculus dextrus, OS: Oculus sinister), (*BCL2*: B-cell lymphoma 2*, BAX*: Bcl-2 Associated X-protein*, CASP3*: Caspase 3*, CASP7*: Caspase 7*, GFAP*: Glial fibrillary acidic protein*, GSS*: Glutathione synthetase*, AIF1*: Allograft inflammatory factor 1*, TLR4:* Toll-like receptor 4, (*p*>0.05, ns= Non-significant) (OD: N=6, OS: N=6).

**Fig. 3.**
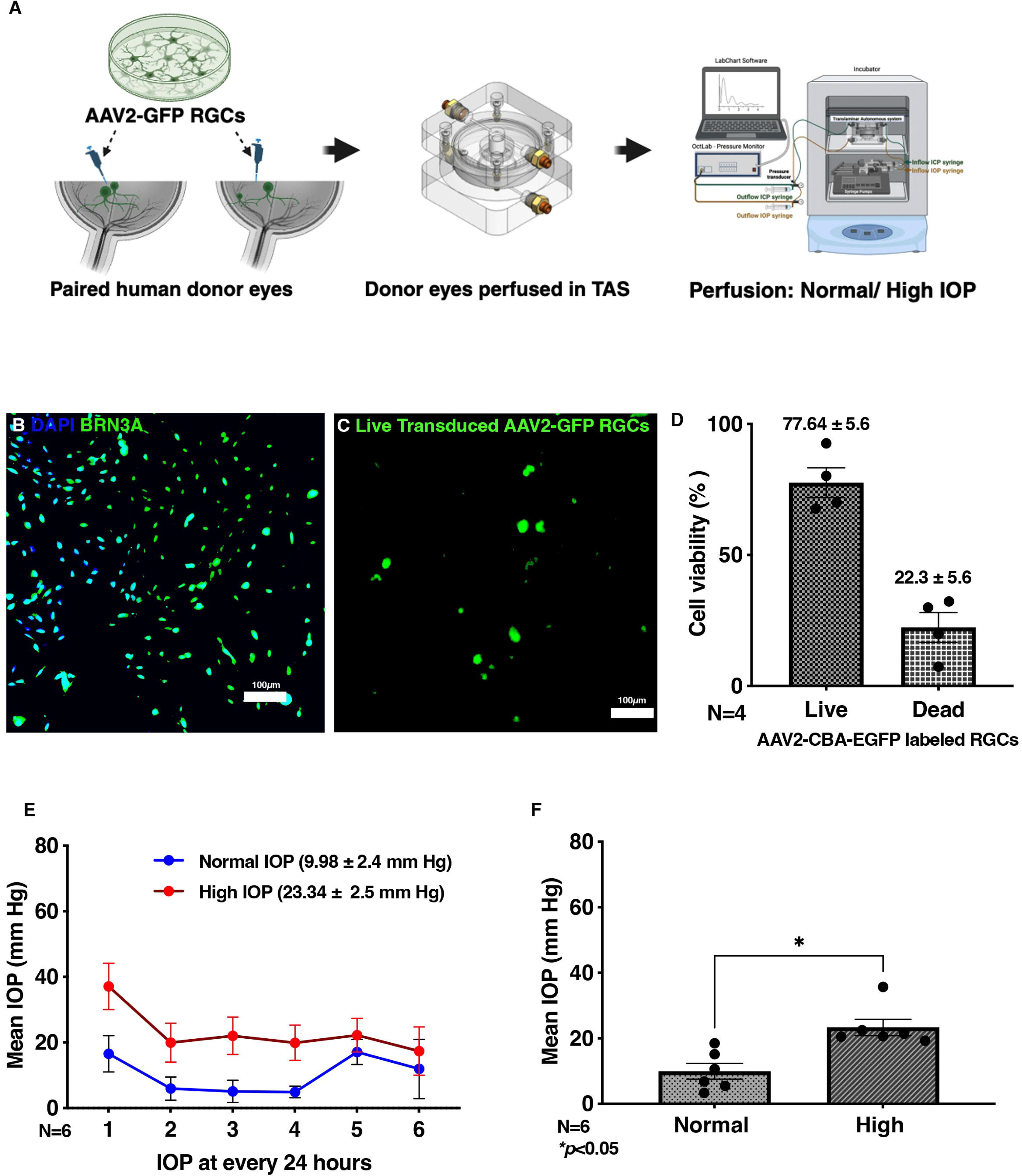
Transduction of RGCs with AAV2-CBA-EGFP and pressure perfusion in the donor eyes after transplantation. A) Illustrating the experimental paradigm for seeding and perfusing human iPSC derived RGCs into human posterior segments. Cultured iPSC derived RGCs depicted positive expression for (B) marker BRN3A and (C) AAV2-CBA-EGFP after 72 hours post-transduction. D) Graphs depicting viability of the AAV2-CBA-EGFP transduced cells before transplantation, (E) graphs depict perfusion pressures used for the experiment over a 6-day interval. F) Graph represents mean IOP over 6 days. iPSC: Induced pluripotent stem cells, IOP: Intraocular pressure, Magnification 20X, *\*p<*0.05.

After a 24-hour post-seeding period, the transplanted eyes were cultured within the TAS model and perfused under normal and elevated intraocular pressure (IOP) conditions for a total of 6 days. As depicted in Fig. 3E, the IOP was successfully maintained at both normal and high conditions over the 6-day period, with significant differences observed between the paired donor eyes (normal IOP: 9.98 ± 2.4 mm Hg; high IOP: 23.34 ± 2.5 mm Hg, \**p<*0.05) (Fig. 3F). Additionally, we would like to note that the normal IOP on day 5 was slightly elevated compared to the other perfusion days. However, it did not exceed pathological levels (Mean normal IOP on day 5 = 17.13 mmHg, N = 6).

### 3.4. Survival of transplanted RGCs in the host human donor retina

Survival of both transplanted and native RGCs in the host retina following pressure perfusion culture was evaluated using gene expression analysis and flat mount immunostaining. We assessed the expression of apoptotic (*BAX, CASP3*), and RGC (*THY1*) markers. As shown in Fig. 4A, the RGC transplanted eyes at high IOP shown a significant increase in *BAX* (normal IOP: 0.89 ± 0.38 FC, high IOP: 4.1 ± 0.64 FC, *\*p<*0.05), and a trend towards increased expression of *CASP3* (normal IOP: 1.6 ± 1.03 FC, high IOP: 1.97 ± 0.69 FC, *p>*0.05, ns) compared to RGC transplanted eyes perfused at normal IOP. Corresponding to these findings, we observed a significant downregulation of *THY1*, a key marker of RGCs, in transplanted eyes subjected to high IOP as depicted in Fig. 4B (normal IOP: 1.29 ± 0.43 FC, high IOP: 0.43 ± 0.13 FC, *\*p*<0.05).

**Fig. 4:**
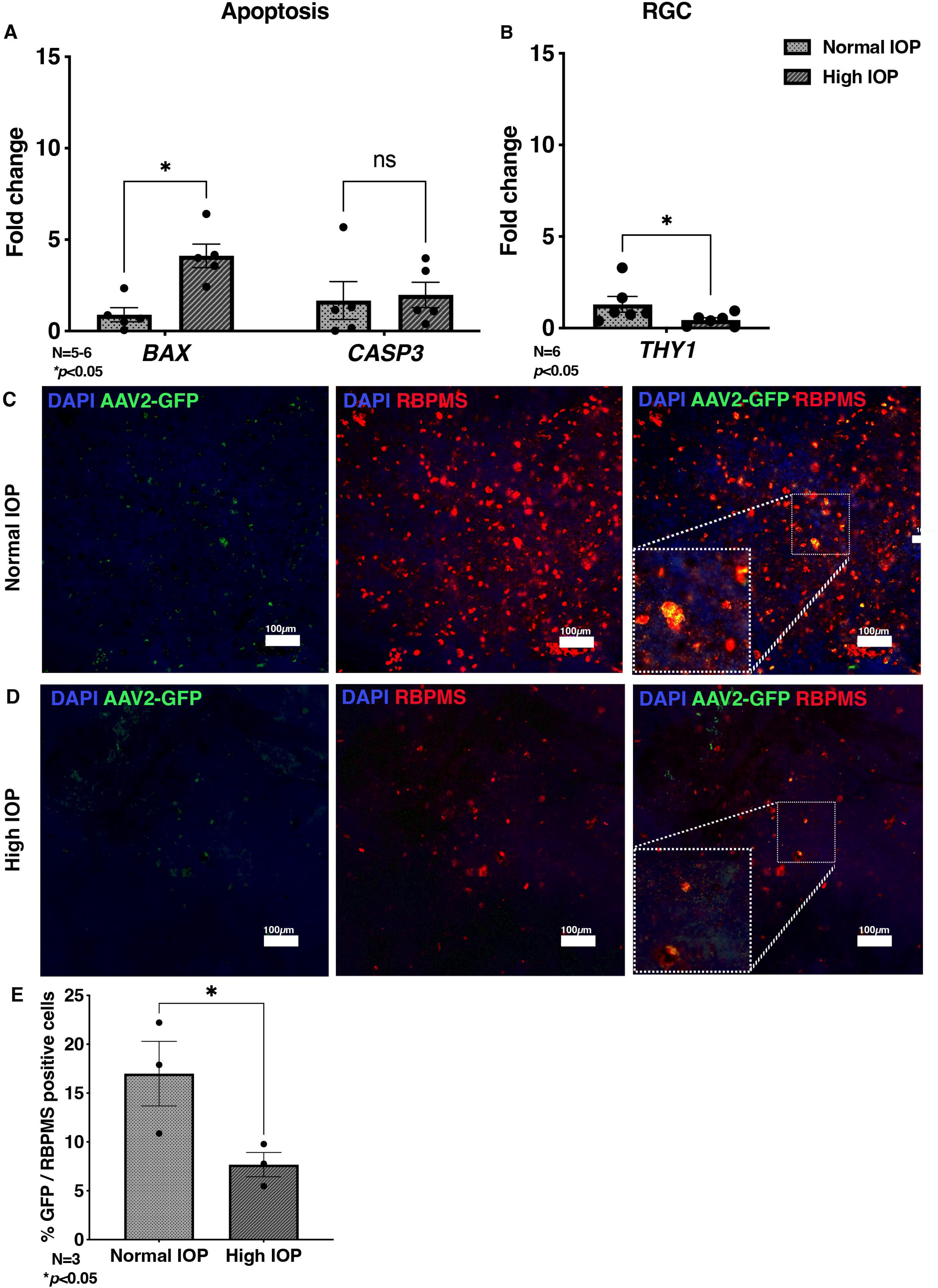
Assessment of apoptosis and RGC survival post transplantation in human donor eyes. Quantitative gene expression of (A) apoptotic (*BAX, CASP3) and* (B) RGC (*THY1*) markers. Flat mount immunostaining of retinal tissues depicting the RGC marker RBPMS and AAV2-CBA-EGFP expression following 6 days of pressure perfusion at (C) normal IOP and (D) high IOP. E) Bar graph depicting the percentage of GFP positive cells over the total number of RGCs. *BAX:* Bcl-2 associated X protein *CASP3*: Caspase 3, *THY1*: Thymus cell antigen 1, IOP: Intraocular pressure, \**p<*0.05, ns = non-significant (N=4), Magnification 20X.

The loss of RGCs transplanted eyes under high IOP was further validated through flat-mount immunostaining of central retinal sections using RBPMS, an RGC specific marker. As depicted in Fig. 4C, the number of native RGCs and AAV2-CBA-EGFP transplanted RGCs was higher under normal IOP conditions. In contrast, transplanted eyes perfused at under high IOP conditions, the survival of both AAV2-CBA-EGFP transplanted RGCs and native RGCs were noticeably compromised, confirming high IOP negatively impacted the survival of native and transplanted RGCs in the human retina. Furthermore, we quantified and calculated the percentage of GFP positive transduced RGCs in normal and high IOP condition over the total number of RBPMS positive RGCs. Consistent with our immunostaining images, we identified a significantly increased percentage of GFP positive cells in the eyes exposed to normal IOP conditions (Normal IOP: 16.9± 3.3%, High IOP 7.6 ± 1.2% N=3, *p<0.05) (Fig.4D).

### 3.5. Increased ECM deposition within transplanted human donor eyes at high IOP compared to normal IOP

To assess whether high IOP induces fibrosis and disrupts the organization of the optic nerve head in RGC transplanted eyes, at post-perfusion we collected optic nerve heads from transplanted eyes subjected to normal and high IOP. The optic nerve heads were stained for ECM markers, such as collagen IV, fibronectin, and laminin. As shown in Fig. 5A, compared to RGC transplanted eyes perfused under normal IOP conditions, those subjected to high IOP perfusion displayed a noticeable increase in expression of these ECM markers post transplantation (Fig. 5B). Further, the mean fluorescent intensity of these ECM markers were quantified after subtracting the background signals (Supplementary Fig. 3), which showed an increased deposition of Collagen IV (Normal IOP: 29 ± 2.9, High IOP: 43.8 ± 10.4 p>0.05,n.s), laminin (Normal IOP: 36.6 ± 11.3, High IOP: 73.6 ± 38.1 p>0.05,n.s), and significant increase in Fibronectin (Normal IOP: 13.6 ± 3.1, High IOP: 196.7 ± 94.06 p<0.05) levels in the transplanted eyes perfused at high IOP conditions (Fig. 5C). These indicate an association of ECM deposition at high IOP condition, which may have an impact on survivability of native RGCs and integration of transplanted RGCs.

**Fig. 5:**
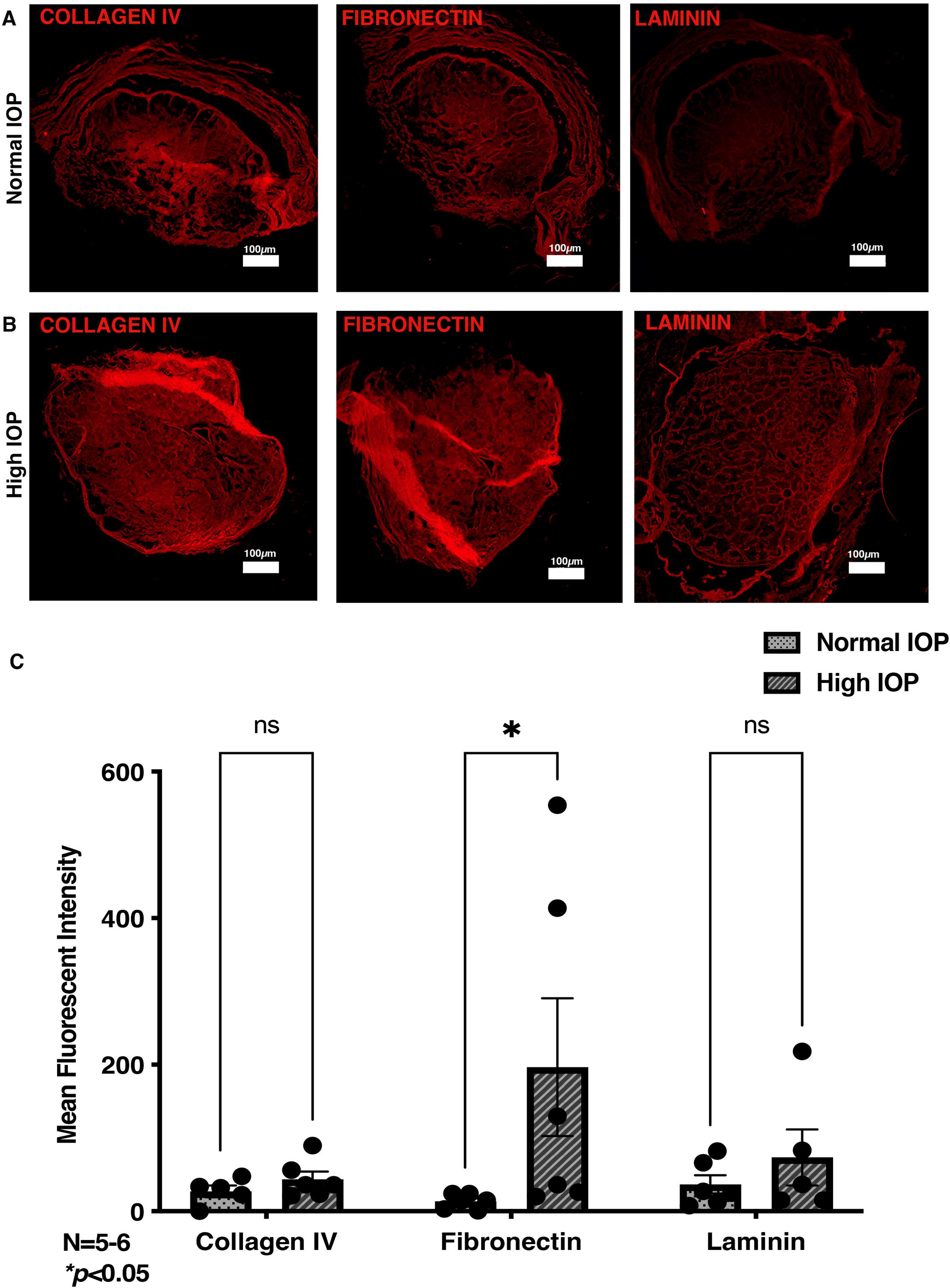
Measurement of ECM deposition in transplanted eyes. Representative images of the optic nerve head depicting the expression of ECM markers such as collagen IV, fibronectin, and laminin under (A) normal IOP compared to (B) high IOP. The mean fluorescent intensity of collagen IV, fibronectin, and laminin calculated through image J quantification. ECM: Extracellular matrix, IOP: Intraocular pressure, ERG: Electroretinography, ns: non-significant, *\*p*<0.05, Magnification 4X.

## 4. DISCUSSION

There is a current need to better characterize RGC transplantation potential as a therapeutic for glaucoma, as retinal pigment epithelial cell transplantation has become an exciting development in clinical trial therapy for other retinal degenerative diseases (Klymenko *et al*., 2024). We found that after transplantation in *ex vivo* posterior human eye cups, purified hiPSC-derived RGCs survived at lower rates under high IOP environment compared to the normal IOP environment given baseline survival rates pre-integration. Our findings show downregulated gene expression of RGC marker, *THY1*, after 6 days of pressurization in high IOP condition eyes compared to normal IOP eyes (Normal IOP: 1.29 ± 0.43 FC, High IOP: 0.43 ± 0.13 FC, *\*p*<0.05).

Additionally, the high IOP conditions had significantly increased expression of apoptotic marker, *BAX*, and higher trends of apoptotic marker *CASP3* was observed. It is also documented that BAX is required for RGC death in inherited glaucoma and in optic nerve crush injury glaucoma models (Libby *et al*., 2005). To delineate the source of inflammation and apoptosis in the transplanted eyes, we compared the expression of apoptotic and inflammatory genes between non-transplanted eyes perfused at normal IOP with transplanted eyes at normal IOP. Similarly, we analyzed gene expression between non-transplanted eyes perfused at high IOP with transplanted eyes at high IOP. The data suggest that under normal IOP conditions, apoptosis and inflammation are contributed equally by both host and transplanted cells. However, under high IOP conditions, there is an increasing trend in apoptotic and inflammatory gene expression, primarily originating from host cells (Supplementary Fig. 4). Our lab has previously maintained healthy posterior eye cup tissue within the TAS model for 30 days and successfully modulated high IOP conditions similar to a diseased glaucomatous environment (Sharma *et al*., 2020). Within the high IOP environment, we could expect to see higher expression of apoptotic markers consistent with findings from previous experiments (Hameed *et al*., 2024b). Although our culture system supports culturing of donor eyes up to 30 days, we limited our experiments to 6 days, based on previous ex vivo transplantation studies (Croteau *et al*., 2022; Zhang and Johnson, 2022) and the observed accelerated rate of RGC death under high IOP conditions for both native and transplanted RGCs by the 6 day time point. Given the decrease in both transplanted RGCs and native RGCs on flat mount staining and gene expression for high compared to normal IOP, this data suggests that the diseased environment (high IOP) was less receptive to transplanted neurons compared to the undamaged retina model (normal IOP).

To assess the transplantation potential of donor RGCs, we employed iPSC-derived RGCs generated from donor keratocytes (Hameed and Sharma, 2025). To ensure the culture was enriched exclusively with RGCs, we selected day 30 ROs for RGC culture, based on the fact that RGCs are the first neural cell type to emerge in ROs typically between days 27 and 33, whereas other neural cell types, such as photoreceptors, develop after this period (Nguyen-Ba-Charvet and Rebsam, 2020). Quantitative RGC survival studies are limited in *ex vivo* human donor eyes. Our study shows successful transplantation of AAV2-CBA-EGFP labeled RGCs within the human host retina. Since AAV2-based transduction can induce inflammation (Gao *et al*., 2009), we evaluated inflammatory markers such as IL-6 and TNF-α in transduced RGCs and compared them with non-transduced cells. Our data demonstrated that transduction did not trigger amplified inflammation in RGCs (Supplementary Fig. 2). Another potential concern is the possibility of AAV2 transmission or cell transfer after transduction. However, since both our normal and high IOP conditions utilized AAV2-transduced cells we can effectively state that both conditions would have this effect and the effects we observe from transduction would not be impacted due to this phenomenon. *In viv*o rodent and primate recipients have survival of transplanted typically at 1-5% with majority of retained transplanted cells remaining healthy after more than a month of transplantation (Venugopalan *et al*., 2016). However, given the limited number of surviving transplanted RGC, these additional neurons may not provide functional replacement of degenerated ganglion cells as seen with the variable outcomes and success in prior findings (Oswald *et al*., 2021; Suen *et al*., 2019; Hertz *et al*., 2014; Jagatha *et al*., 2009). Further, recent studies have focused on a secondary benefit of transplanted RGCs, which may also provide neuroprotection in disease models of RGC death (Mead *et al*., 2013; Singhal *et al*., 2012; Cho *et al*., 2012). A study of human stem cell derived RGCs intravitreally transplanted in *in vivo* mice found protective effects from neurodegeneration after optic nerve trauma (Luo *et al*., 2022). A proposed mechanism for the protective effect of transplanted RGCs is the transference of extracellular vesicles containing diverse, multifactorial cargo to endogenous RGCs that may play a pivotal role in cellular health (Lucci and De Groef, 2023).

Further, there are estimated to be over 30 subtypes of RGCs, each with unique molecular and morphological signatures (Baden *et al*., 2016). Future studies can also examine whether certain subtypes of RGCs would be more receptive to integration after transplantation. Previous data has shown alpha-RGCs and intrinsically photosensitive RGCs are less sensitive to cell death than other RGC subtypes in a mouse model of glaucoma (Struebing *et al*., 2016). When looking at the different gene marker expression for RGCs types, only *THY1* was significantly lower in the high IOP setting, suggesting generalized decrease of transplanted RGCs to seeding or survival in diseased environment compared to the normal conditions. The poor integration of RGCs in high IOP conditions can be partially explained by the increased expression of ECM markers seen in high IOP conditions in our optic nerve head cross-section staining experiment. The high IOP promoted fibrosis and ECM disorganization (Fig. 4), preventing survival of transplanted RGCs and inducing further damage to native RGCs.

The clinical relevance of our findings demonstrates the differences in RGC transplantation between normal and diseased states suggest that transplantation as a therapeutic strategy may be more beneficial during early stages of glaucoma and only after the IOP is controlled with medications. At earlier stages of glaucoma, there is a decreased number of diseased neurons, requiring a lower yield of transplanted RGCs needed to replace them. There is also a higher number of healthy remaining neurons and thus a stronger potential for transplanted RGCs to confer early protection of the healthy endogenous RGCs (Luo *et al*., 2022). Additionally, we found that higher IOP conditions, as observed in later and more severe stages of glaucoma, did not allow for transplanted RGCs to successfully survive within the host retina. Given that high IOP creates a diseased microenvironment for RGCs, future studies should explore whether prior high IOP conditions can affect transplanted RGC survival even when transplantation occurs under a normal IOP environment. This would help clarify whether cell survivability is directly influenced by high IOP or by the microenvironment, providing insight into the potential benefits of this therapy for IOP-controlled glaucoma.

A limitation of our study acknowledges the lack of direct imaging for visualization of transplanted RGC’s migration and integration into the host ganglion cell layer and formation of intercellular connections. This has previously been shown in similarly transplanted GFP-positive RGCs, which indicated migration into the ganglion cell layer, new growth cone-like structures from transplanted RGCs, and synaptic formation between transplanted RGCs and host retina (Venugopalan *et al*., 2016). While our study does not directly evidence synaptic formation with synaptic marker staining, we measured electrophysiological activity by utilizing OcuScience *ex-vivo* ERG system of the explanted host retina. The changes in electrophysiological activity demonstrated a decreased b/a ratio with increase in implicit times for both a-wave and b-wave at 30 mcd flashlight intensity in high IOP perfused eyes. This shows a negative trend in overall retinal activity in the diseased state despite treatment of transplanted RGCs (Supplementary Fig. 5). However, the functional activity of the retina needs to be evaluated using larger sample size in future studies along with assessment of functional integration of transplanted cells into host retina.

Additionally, our study used axotomized human posterior eye cups which have a basal level of degeneration occurring compared to an *in vivo* model. To control for the higher level of neural degeneration than a healthy perfused eye, the eyes were normalized to data sectioned from contralateral retina before the start of perfusion. Our previous studies have shown maintenance of retinal tissue integrity, structure, and complexity under conditions modulated by the TAS model similar in this study (Sharma *et al*., 2020). This ensures that any previous degeneration before starting the experiment was normalized to tissue that has undergone the same timeframe of axotomization and neural degeneration. The neural degeneration in the axotomized eye may model the glial fibrosis at the optic nerve head in glaucoma, secondary to increased IOP on the lamina cribosa. The elevated IOP causes RGC axonal transport deficits and neurotropic factor delivery impairment (Claes *et al*., 2019), leading to RGC degeneration and death (Burgoyne, 2011). While glaucomatous cell loss occurs over time, our axotomized *ex vivo* model provides an acute model of a chronic process. Given that our findings suggest an improvement and maintenance of retina cell health in a basally degenerating retina, our research shows great potential of using RGC transplantation therapy in living human eyes.

A limitation of the TAS model is the lack of vascular pressure modulation, due to lack of blood circulation within the donor eye. However, the model’s ability to regulate intraocular pressure allows us to specifically delineate the pathogenic effects of IOP changes on transplanted RGC survivability. Another limitation of the TAS model is the inability to model the cyclic circadian rhythms of IOP and ICP observed in *in vivo* human eyes, which may impact RGC survivability under normal physiologic conditions. Another important concern was the IOP fluctuations within the TAS model over time, which could influence RGC survival. However, since these fluctuations occurred in both normal and high IOP-perfused eyes, we attribute our findings primarily to the consistently elevated IOP maintained throughout the 6-day period (23.34 ± 2.5 mmHg), which was significantly higher than the normal IOP treatment group (9.98 ± 2.4 mmHg). Future studies may examine how various RGC subtypes survive in normal and high IOP environments using gene expression biomarkers to quantify survival rates. It would also be valuable to examine differences in RGC survival between transplanted and non-transplanted eyes with cross section optic nerve staining and quantifyingwhether there is a neuroprotective property of transplanted RGCs in the human retina. The studies can also examine improvements in electrophysiologic response to light and measurements of neurite outgrowth and orientation. Overall, our studies show poor outcomes of RGC transplantation in a high IOP state, and many more questions need to be addressed regarding the therapeutic application.

## 5. CONCLUSION

Patients affected by the irreversibly progressive course of glaucoma could benefit from therapeutic options for vision recovery. Utilizing RGC replacement therapy derived from autologous stem cells would be a valuable option. Our data show that RGCs differentiated from hiPSCs when exposed to normal IOP in *ex vivo* donor eyes have improved survival outcomes, and less fibrosis compared to those under elevated IOP. This suggests that stem cell therapies in glaucoma might improve outcomes when used alongside current IOP-lowering therapies at earlier and controlled stages of glaucoma. Our study adds to the growing body of knowledge for stem cell therapy for tissue replacement in glaucomatous retinas and represents a proof of concept for their transplantation into *ex vivo* donor eyes.

IOP: Intraocular pressure
RGC: Retinal ganglion cell
hiPSC: Human induced pluripotent stem cell
TAS: Translaminar autonomous system
ERG: electroretinography
RO: Retinal organoids
ECM: Extracellular matrix
FC: Fold change
AAV2-CBA-EGFP: Adeno-Associated Virus serotype 2- Chicken Beta Actin- Enhanced Green Fluorescent Protein

## FUNDING

This work was supported by an unrestricted grant from Research to Prevent Blindness, Inc. to the Indiana University School of Medicine, Department of Ophthalmology. The work was also supported by the Eugene and Marilyn Glick Eye Institute Startup Funds and grant from National Institute of Health (1R01EY034174).

## AVAILABILITY OF DATA AND MATERIALS

All the data generated and analyzed during this study have been included in this published article and its supplementary information files.

## CREDIT AUTHORSHIP CONTRIBUTION STATEMENT

SSH together with SJG performed the design and writing for the original draft of the manuscript. SSH was responsible for formal analysis of the data, performed analysis, investigation, data acquisition for the manuscript. KKH drafted the discussion and performed critical review of the manuscript. TPS was responsible for the conceptualization, project design, funding acquisition, project administration, supervision, reviewing and approving the final version of the manuscript.

## COMPETING INTEREST

The corresponding author declares a conflict of interest on the TAS model, for which she holds US11678658B2.

