## Supplemental Figure for "Survival of Transplanted Retinal Ganglion Cell in Human Donor Eyes under Elevated Pressure"

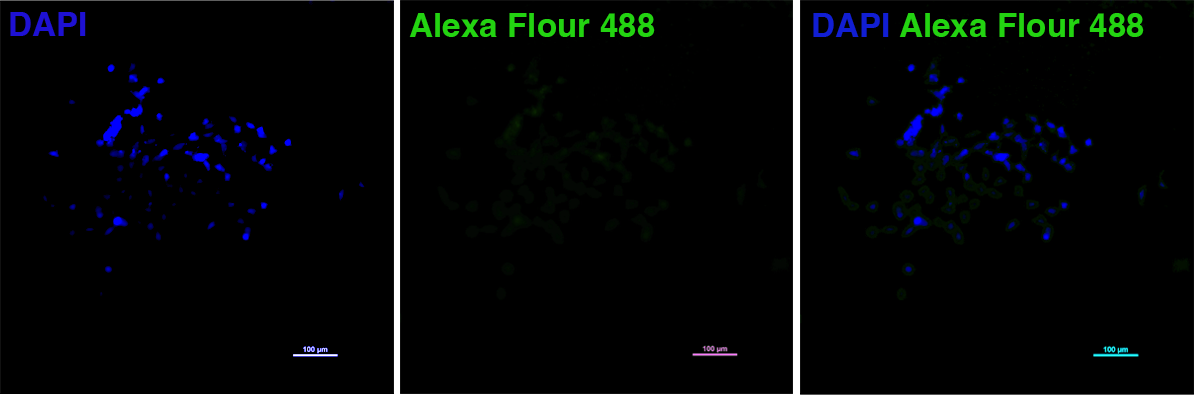

**Supplementary Fig. 1.** The secondary antibody control staining showing minimal non-specific binding of secondary antibody to the dissociated RGCs, confirming BRN3A specificity.

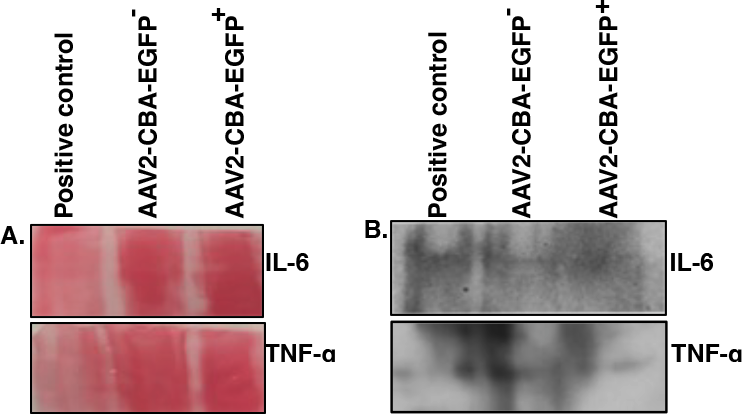

**Supplementary Fig. 2.** Assessment of inflammation In AAV2-CBA-EGFP transduced RGCs as compared to non-transduced RGCs. (A) Ponceau S staining of the proteins (B) western blotting images showing the expression of IL-6 and TNF-α. The conditioned medium collected from high IOP eyes were used as a positive control.

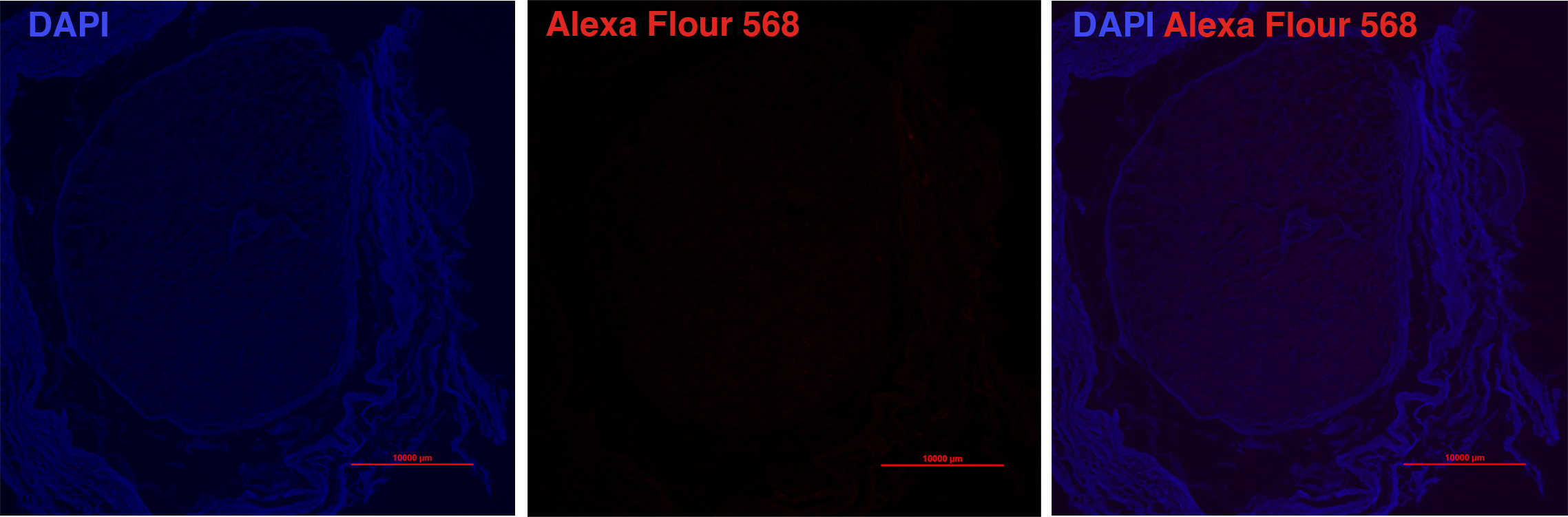

**Supplementary Fig. 3.** The secondary antibody control staining showing no non-specific binding of secondary antibody to the optic nerves.

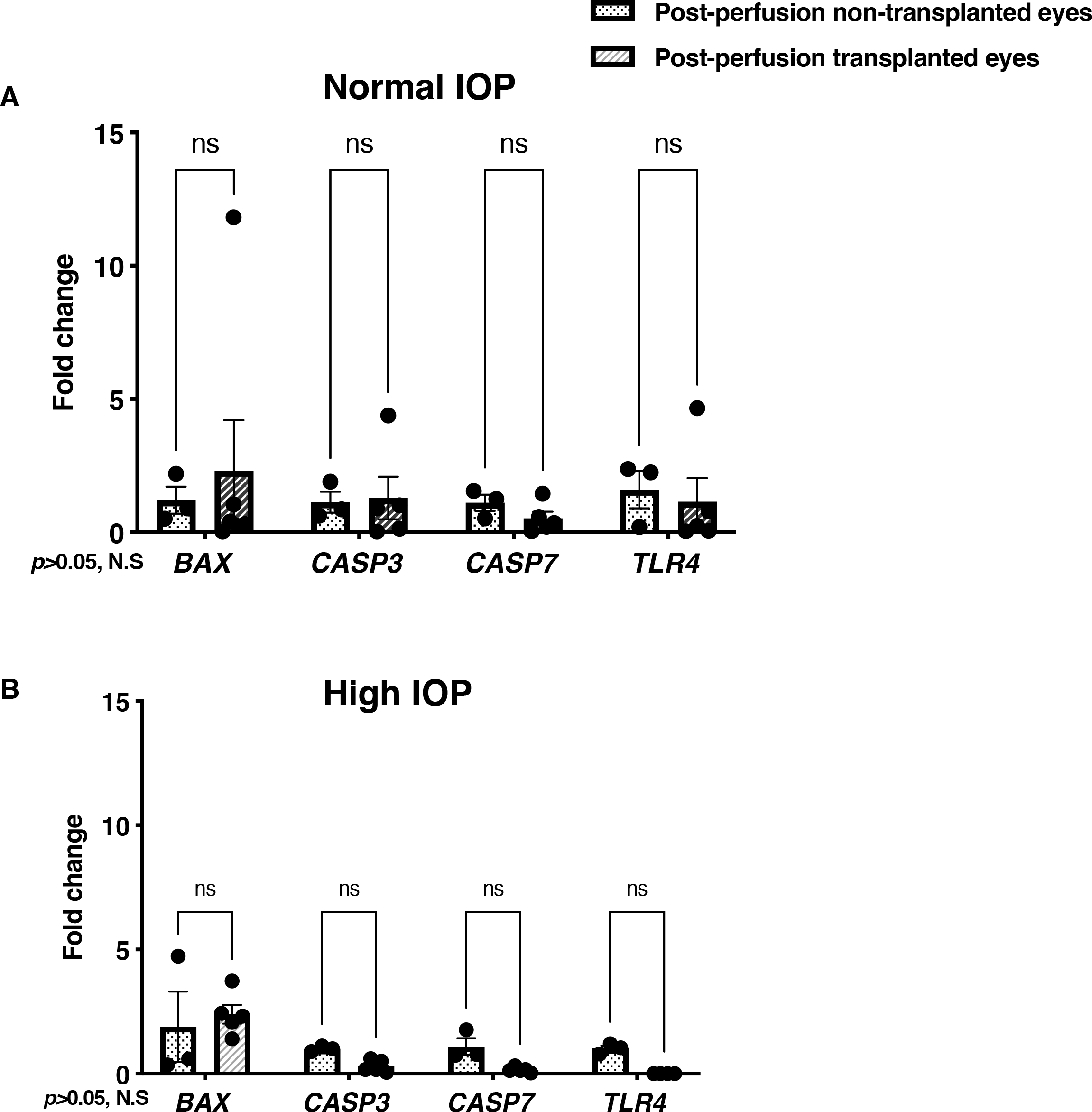

**Supplementary Fig. 4.** The qPCR data presents the quantitative gene expression of apoptotic and inflammatory markers in perfused eyes under two conditions: (A) normal IOP, compared to normal IOP eyes with transplanted RGCs, and (B) high IOP, compared to high IOP eyes with transplanted RGCs. Sample sizes include N=3 for non-transplanted eyes at both normal and high IOP, and N=6 for eyes with RGC transplantation. *BAX*: Bcl-2 Associated X-protein*, CASP3*: Caspase 3*, CASP7*: Caspase 7*,* *GFAP*: Glial fibrillary acidic protein*, TLR4:* Toll-like receptor 4, ns: non-significant.

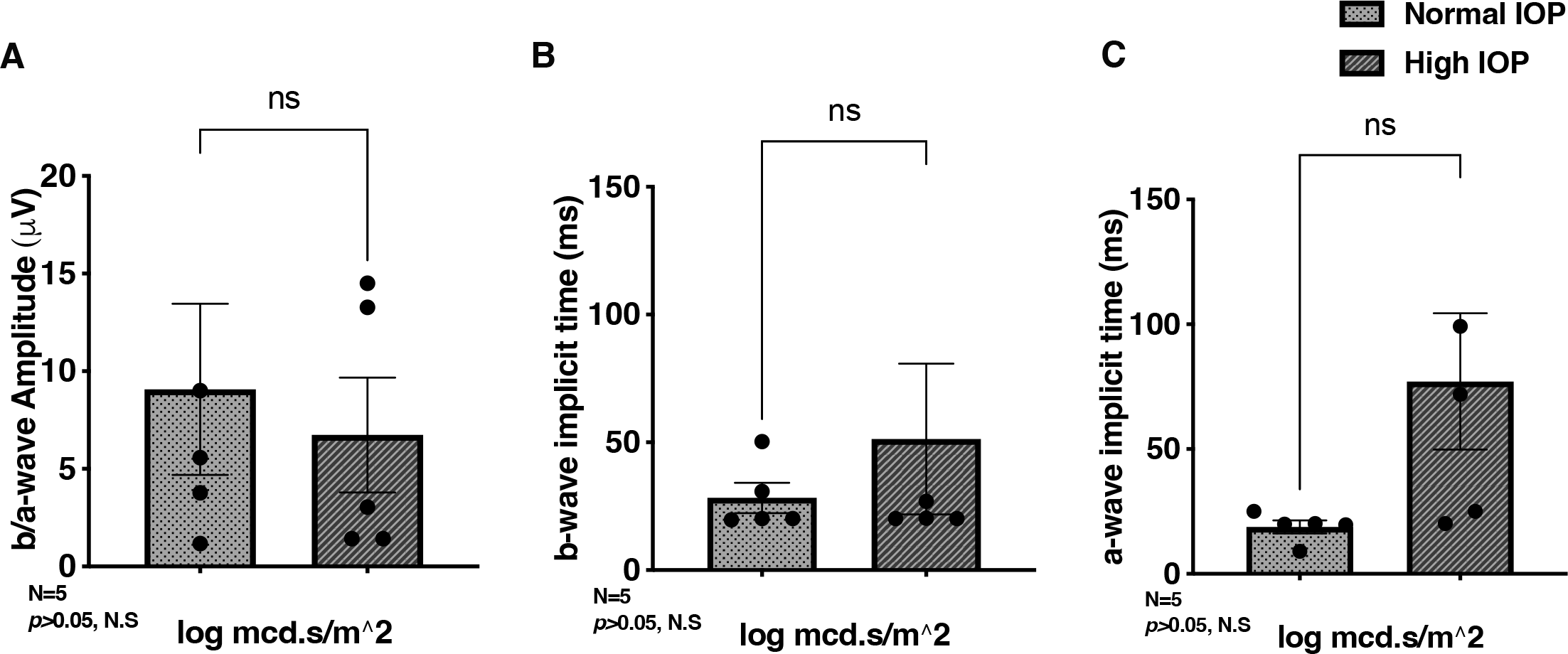

**Supplementary Fig. 5.** Measurement of retinal activity through *ex vivo* ERG. The *ex vivo* scotopic ERG performed on retinas post-perfusion indicated a trend toward decreased (C) b/a ratio, along with higher implicit times of both (D) a-wave and (E) b-wave at high IOP compared to normal IOP. ECM: Extracellular matrix, IOP: Intraocular pressure, ERG: Electroretinography, ns: non-significant, mcd: millicandela, ms: millisecond, µV: microvolt, ns: non-significant.
